# Acylation of the RTX Toxin MbxA stimulates host membrane disruption through a specific interaction with cholesterol

**DOI:** 10.1101/2025.02.19.639210

**Authors:** Feby Mariam Chacko, Sarah Michelle Ganz, Anne Pfitzer-Bilsing, Sebastian Hänsch, Philipp Westhoff, Stefanie Weidtkamp-Peters, Sander H. J. Smits, Marten Exterkate, Lutz Schmitt

## Abstract

RTX toxins, characterized by their calcium-binding glycine-rich repeat regions, are known for their ability to disrupt host cell membranes. Although the acylation of specific lysine residues in these toxins is established as crucial for their hemolytic activity, the precise mechanisms underlying this enhancement remain unclear. In this study, we explore the role of acylation in the pore-forming behaviour of the RTX toxin MbxA by comparing the lytic activities of acylated MbxA and its non-acylated form, proMbxA, on *in vitro* membrane systems, as well as living cells. Our findings demonstrate that a cholesterol specific interaction promotes MbxA- induced membrane disruption in an acylation-dependent manner. Further analysis revealed that acylation is not necessary for initial membrane binding but markedly enhances pore formation. Overall, our results offer valuable insights into the molecular determinants that regulate RTX toxin activity, highlighting a complex interplay between lipid composition (sterols), acylation, and membrane disruption, thereby advancing our understanding of RTX toxin pathogenesis.

## Introduction

RTX (repeats-in-toxin) toxins are a family of exoproteins secreted by Gram-negative bacteria through a Type-1 secretion system (T1SS) [1]. Many bacteria utilize these pore-forming RTX toxins as virulence factors during infection. For instance, the uropathogenic *Escherichia coli* (UPEC) hemolysin A (HlyA) is an RTX toxin linked to urinary tract infections (UTIs), one of the most common bacterial infections worldwide [2]. Another example is the Adenylate Cyclase Toxin (ACT) from *Bordetella pertussis*, which targets the human respiratory tract and causes whooping cough, a highly contagious and recurring disease [3]. The oral bacterium *Aggregatibacter actinomycetemcomitans*, known to cause localized aggressive periodontitis as well as systemic infections such as endocarditis, produces an RTX protein called leukotoxin A (LtxA), which is lethal to human immune cells [4].

RTX proteins are distinguished by the presence of glycine-rich nonapeptide repeats (GGxGxDxUx; where x represents any amino acid and U indicates a large hydrophobic amino acid) located in the C-terminal part of the protein [5]. HlyA serves as the prototype of the RTX toxin family. In the cytoplasm of the bacteria, the translated HlyA remains unfolded and inactive prior to secretion. It becomes active through post-translational acylation at two lysine residues, K564 and K690, which is catalyzed by the acyl transferase HlyC. The acylated HlyA is then secreted via the T1SS, an ABC transporter composed of HlyB (the inner membrane protein), HlyD (the membrane fusion protein), and TolC (the outer membrane protein). The binding of the C-terminal secretion signal of HlyA to HlyB and/or HlyD recruits TolC to the outer membrane, forming a complete channel for transporting the unfolded toxin to the extracellular space. The C-terminal region of HlyA is the first to reach the extracellular environment. The higher concentration of Ca^2+^ ions outside the bacteria compared to the cytoplasm promotes the binding of these ions to the RTX domain of the secreted HlyA, triggering the protein’s folding. Once the entire HlyA protein is outside the cell, it fully folds and becomes active, ready to form pores in various types of cells [6, 7].

Secreted RTX toxins generally exhibit a broad range of host cell specificity [5, 8]. Many RTX toxins have been shown to interact with specific β2 integrin receptors located on the cell membranes of certain host cells, such as Jurkat cells [2, 9–18]. Removal of these β2 integrin receptors from host cell membranes, or blocking them with antibodies, has been associated with reduced susceptibility of the host cells to RTX toxin-induced cytotoxicity [12–15]. However, RTX toxins can also induce cytotoxicity in cells that lack β2 integrin receptors, such as red blood cells (RBCs) [19, 20]. Furthermore, the pore-forming and lytic properties of RTX toxins have been demonstrated in various artificial membranes, including liposomes, Giant Unilamellar Vesicles (GUVs), and supported lipid bilayers (SLBs) [21, 22]. Not all RTX toxins rely on β2 integrin receptors to interact with host cell membranes. For instance, the RTX toxin RtxA from *Kingella kingae* does not engage with β2 integrin receptors at all [23]. These findings suggest a potential β2 integrin receptor-independent mechanism for RTX toxin interaction with host cell membranes. Several RTX toxins have been found to associate with non-protein structures on host cell membranes, such as the carbohydrate chains of glycophorins, gangliosides, and cholesterol, supporting the idea of a receptor-independent interaction pathway [3, 23–33]. While these host cell surface structures are known to interact with RTX toxins, the precise mechanism by which they facilitate RTX toxin pore formation remains poorly understood.

*Moraxella bovis* is an intracellular parasite that inhabits mucous membranes and causes the ocular disease infectious bovine keratoconjunctivitis (IBK) in cattle. MbxA is a rather uncharacterized RTX toxin produced by *M. bovis*. In this study, we employed a heterologous system to secrete both the acylated, active form of MbxA and the non-acylated, inactive proMbxA using a T1SS in *E. coli*, specifically the HlyA secretion system [34]. MbxA shares 42% sequence identity with HlyA, and the RTX operon of *M. bovis* contains all the genes required for activation (*mbxC*) and secretion (*mbxB*, *mbxD*, and *tolC*) of an RTX toxin [34–41].

Our previous work demonstrated that these heterologously secreted and purified MbxA proteins could induce cytotoxicity in human epithelial (HEp-2) cells, T cells, and sheep red blood cells, whereas proMbxA remained inactive against these cell types [34]. Here, we combined an *in vitro* and *in vivo* approach to further explore the broad host cell specificity of MbxA, focusing on β2 integrin receptor-independent interactions. More specifically, we systematically studied the role of the membrane lipid composition, revealing a prominent role for cholesterol in the pore-forming mechanisms of MbxA.

## Results

### The liposome leakage assay reveals a specific lipid dependency in the lytic activity of MbxA and proMbxA

To examine the β2 integrin receptor-independent pore-forming activity of MbxA, we conducted a commonly used liposome leakage assay, based on the fluorophore 8-Aminonaphthalene- 1,3,6-Trisulphonic acid (ANTS) and its quencher p-Xylene-Bis-Pyridinium Bromide (DPX) (figure 1a). After establishing baseline fluorescence, 100 nM of either MbxA or proMbxA was added to the liposomes, thereby initiating possible pore formation/leakage. After reaching a stable plateau, 0.5% Triton X-100 was introduced, dissolving all liposomes. As a consequence, the measured fluorescence represents 100% content release (figure 1b).

**Figure 1:**
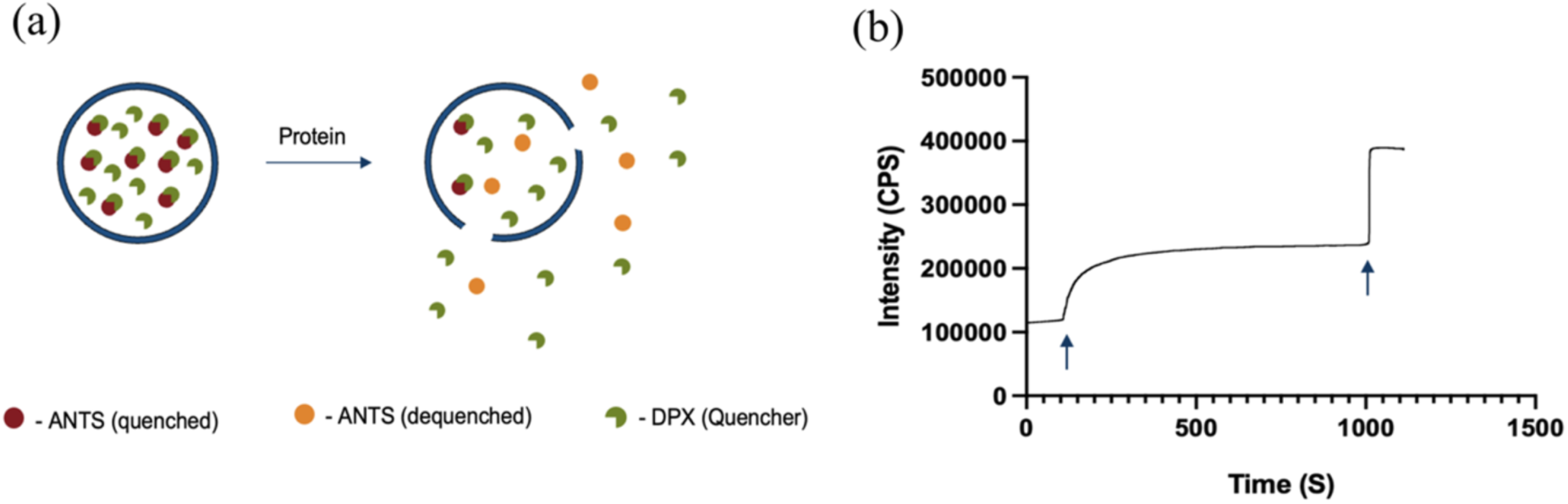
(a) Liposomes with various lipid compositions were encapsulated with the fluorophore ANTS and the quencher DPX. Upon addition of protein (MbxA or proMbxA) pores might be formed, resulting in leakage and consequently dequenching of ANTS. (b) Fluorescence measurement of DOPC liposomes encapsulated with ANTS/DPX prepared from 12 µM DOPC. After 100 sec, 100 nM MbxA is added (first arrow). Once a plateau is reached, 0.5% Triton X-100 was added (second arrow), resulting in the complete release of the liposomal content. ANTS: 8-Aminonaphthalene-1,3,6-Trisulphonic acid, disodium salt; DPX: p-Xylene-Bis-Pyridinium Bromide.

### Increased lytic activity of MbxA and proMbxA for liposomes composed of unsaturated fatty acids and negatively charged lipid headgroups

To investigate the lipid dependency of pore formation by MbxA and proMbxA, we assessed the lytic activity of the proteins on a series of liposomes with varying lipid compositions based on either the saturation of their lipid tails, or the charge of the headgroup. Focusing on the acyl chains three different lipid species were chosen: di-palmitoyl phosphatidylcholine (DPPC) which has two saturated acyl-chains, palmitoyl-oleoyl phosphatidylcholine (POPC) containing one unsaturated acyl chain, and di-oleoyl phosphatidylcholine (DOPC) which consists of two unsaturated acyl chains. Both MbxA and proMbxA exhibited a higher pore-forming efficiency in the presence of the highly unsaturated DOPC liposomes, while only limited pore formation was observed for the fully saturated DPPC liposomes (fig. 2a) as evident by the lower fluorescence increase after protein addition for DPPC liposomes. Notably, MbxA demonstrated a specific activity in the presence of POPC liposomes compared to proMbxA, illustrating that the acyl-moieties of the protein effect the lytic activity under specific membrane conditions. Subsequently, the lytic activity of MbxA and proMbxA was tested in the presence of different head groups, differing in their overall charge. While POPC possesses a net neutral charge, palmitoyl-oleoyl phosphatidylserine (POPS) has a net negative charge. Both MbxA and proMbxA exhibited increased lytic activity toward the negatively charged liposomes (fig. 2b). Notably, the release of liposomal content from the POPC:POPS 1:1 mixture was intermediate between that of the POPC and POPS liposomes for both MbxA and proMbxA, indicating no specific interaction between the acyl-chain moieties of the protein and the lipid headgroup.

**Figure 2:**
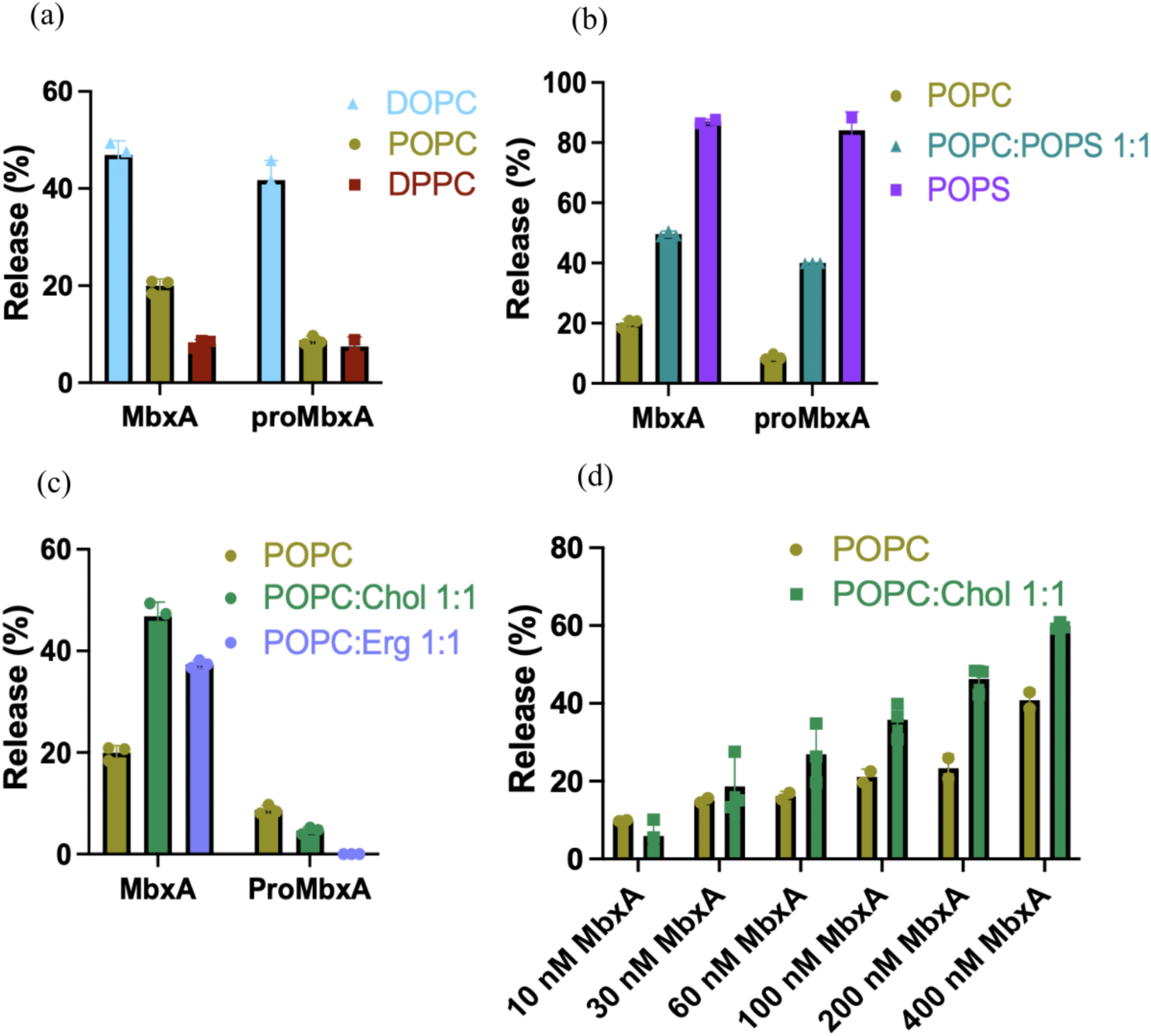
The percentage release of liposomal content by the lytic activity of MbxA and proMbxA on different sets of liposomes. Lytic activity of MbxA and proMbxA on liposomes based on (a) the acyl chain saturation of the lipid chain, (b) the charge of the lipid head group, and (c) cholesterol and ergosterol incorporation into the liposomes. (d) Percentage release of liposomal content when different concentrations of MbxA were incubated with 12 uM POPC and POPC:Chol 1:1 liposomes.

### MbxA exhibits specific lytic activity towards cholesterol-containing liposomes, while proMbxA does not

Given that cholesterol is present in mammalian cell membranes but absent in bacterial cell membranes, we assessed whether MbxA and proMbxA exhibit specific lytic activity against cholesterol-containing membranes. When cholesterol was added to POPC liposomes at a 1:1 ratio, a 2.5-fold increase in liposome leakage was observed for MbxA compared to POPC liposomes, whereas proMbxA showed no such increase (Fig. 2c).

A specific role of the acylation moieties of MbxA in the lytic activity of the protein was already observed for POPC liposomes but seems to be further enhanced in the presence of cholesterol. To further elucidate this phenomenon, we tested the lytic activity of POPC vs. POPC:Chol 1:1 liposomes at varying MbxA concentrations (10 nM, 30 nM, 60 nM, 100 nM, 200 nM, and 400 nM) (Fig. 2d). For both sets of liposomes, we observed a linear increase in the release of liposomal content, but the POPC:Chol 1:1 liposomes exhibit an overall higher release, thereby confirming the stimulating effect of cholesterol on the lytic activity of MbxA.

Furthermore, we examined the lytic activity of MbxA and proMbxA in the presence of sterols other than cholesterol by incorporating ergosterol into POPC liposomes (POPC:Ergosterol 1:1). Remarkably, similar to cholesterol, ergosterol enhanced the leakage of liposomal content when incubated with MbxA, although it did not have the same effect on proMbxA (Fig. 2c).

Given that gangliosides have been reported as receptors for other RTX toxins [26, 27, 31], we further investigated the effect of gangliosides on the lytic activity of MbxA and proMbxA. However, POPC liposomes containing 30% GM1 gangliosides did not induce any significant lytic activity for either MbxA or proMbxA; in fact, the lytic activity of MbxA decreased by approximately 5% compared to that observed with POPC liposomes (Supplementary Fig. 2c).

### MbxA, but not proMbxA, causes rupture of POPC:Chol GUVs

To visualize the different leakage effects of MbxA vs. proMbxA on POPC:Chol 1:1 liposomes, we prepared Giant Unilamellar Vesicles (GUVs), incubated them with 100 nM of either MbxA or proMbxA and analyzed them by microscope (Evos M5000 imaging system) over time. Upon incubation with MbxA, noticeable changes in the shape of the GUVs were observed within 30 seconds (Fig. 3b), but they returned to their normal round shapes within 3 minutes, shown with colored arrow marks (Fig. 3c). Ultimately, after 10 minutes, the incubation with MbxA resulted in complete destabilization of the GUVs (Fig. 3d). In contrast, the incubation of 100 nM proMbxA with GUVs did not induce any disturbances in the GUV architecture, and they remained stable for 30 minutes (Fig. 3f to j).

**Figure 3:**
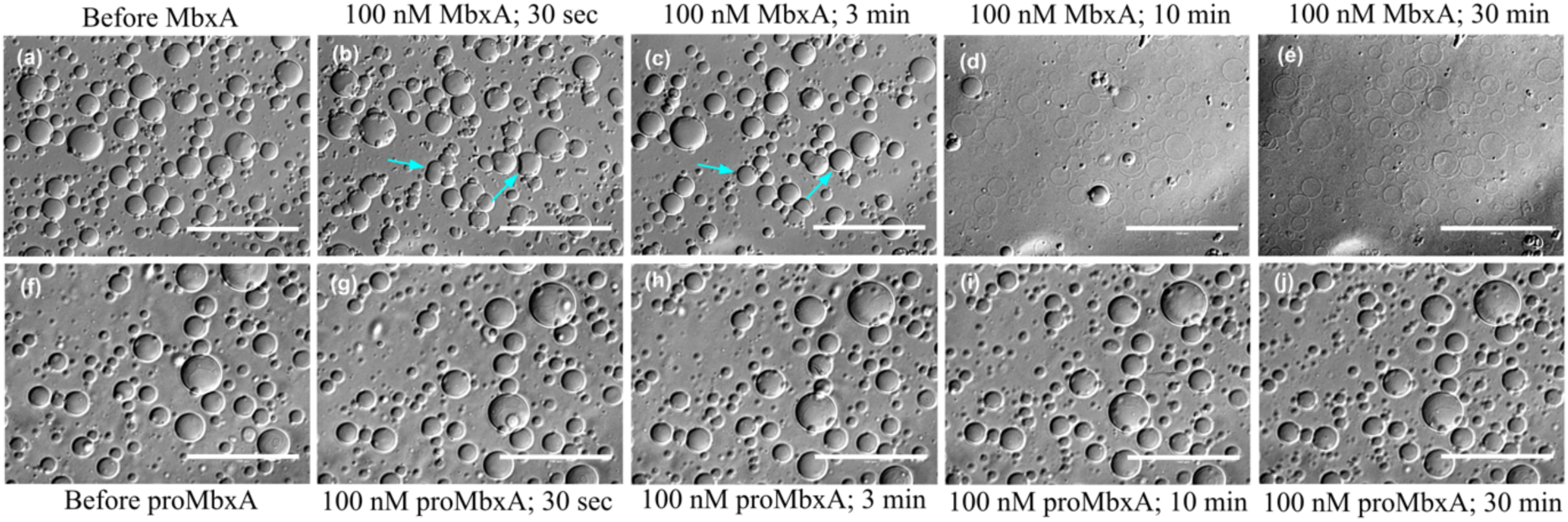
POPC:Chol 1:1 Giant Unilamellar Vesicles (GUVs) (a) before treatment with MbxA, (b) 30 seconds (c) 3 minutes (d) 10 minutes and (e) 30 minutes after incubating with 100 nM MbxA, (f) before treatment with proMbxA, (g) 30 seconds (h) 3 minutes (i) 10 minutes and (j) 30 minutes after incubating with 100 nM proMbxA. Change in GUV shape upon MbxA incubation and its return back to the original shape is shown with colored arrow marks (b&c). Scale bar: 100 μm.

### Flotation assay reveals that acyl chains are not essential for toxin binding to the membrane

Previous experiments (Fig. 2c) indicated that the acyl chains play a crucial role in the lytic activity of the protein within cholesterol-containing membranes. To assess the importance of the acyl chains in the binding of the protein to these membranes, we conducted a flotation assay using MbxA and proMbxA. In this assay, we established a sucrose gradient ranging from 30% at the bottom to 0% at the top. Initially, liposomes incubated with the proteins reside in the bottom fraction. Following ultracentrifugation, the buoyancy of the liposomes causes them to float to the top fraction. If the protein successfully binds to the liposomes, the liposome-bound proteins will also float to the top fraction alongside the liposomes and can be detected via SDS-PAGE (Fig. 4a).

**Figure 4:**
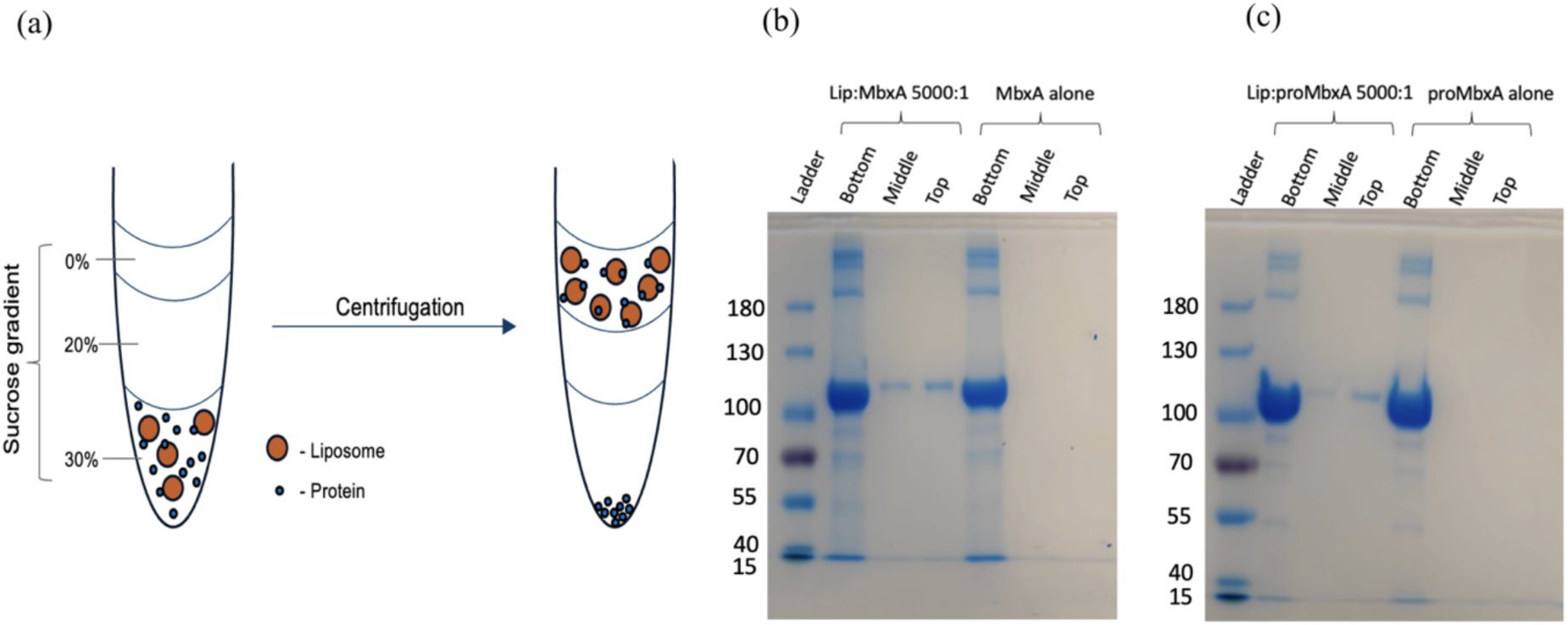
(a) Schematic representation of the flotation assay. (b) Flotation assay with POPC: Chol 1:1 liposome and MbxA (Lipid: MbxA in 5000:1 ratio). (c) Flotation assay with POPC: Chol 1:1 liposome and proMbxA (Lipid: proMbxA in 5000:1 ratio).

The flotation assay using POPC:Chol 1:1 liposomes and MbxA demonstrated the presence of MbxA in all three fractions: bottom, middle, and top (Fig. 4b). Although, most of the protein was found in the bottom fraction, probably aggregated protein, some protein was detected in the middle and top fraction, indicating liposomal binding. Given the definition of the volume for each fraction, it is expected to observe some protein crossover in the middle fraction from the top fraction. Importantly, the control experiment, which included only MbxA protein without any liposomes, revealed no protein in the top and middle fractions (Fig. 4b). Similarly, the flotation assay with POPC:Chol 1:1 liposomes and proMbxA indicated that proMbxA also binds to these liposomes (Fig. 4c), suggesting that the acyl chains are not crucial for the protein for its binding to the membrane.

### AFM reveals cholesterol-dependent insertion of acylated MbxA

To investigate the membrane association and coupled pore formation by MbxA in more detail, we conducted Atomic Force Microscopy (AFM) using Supported Lipid Bilayers (SLBs) on a mica surface. AFM images of SLBs composed of POPC or POPC:Chol 1:1 displayed distinct background stripe pattern before treatment with any proteins (Fig. 5a). Following the incubation of SLBs with 100 nM MbxA or proMbxA, we observed an increase in surface height, indicating binding of the protein to the SLB. Notably, MbxA proteins remained stably bound to the POPC:Chol 1:1 SLBs even after three rounds of rigorous washing of the preincubated SLBs (Fig. 5b). In contrast, proMbxA proteins were entirely removed from the SLBs upon washing with buffer, resulting in the reappearance of the background signal. (Fig. 5c). These results implicate that the binding of MbxA to the membrane is more tight.

**Figure 5:**
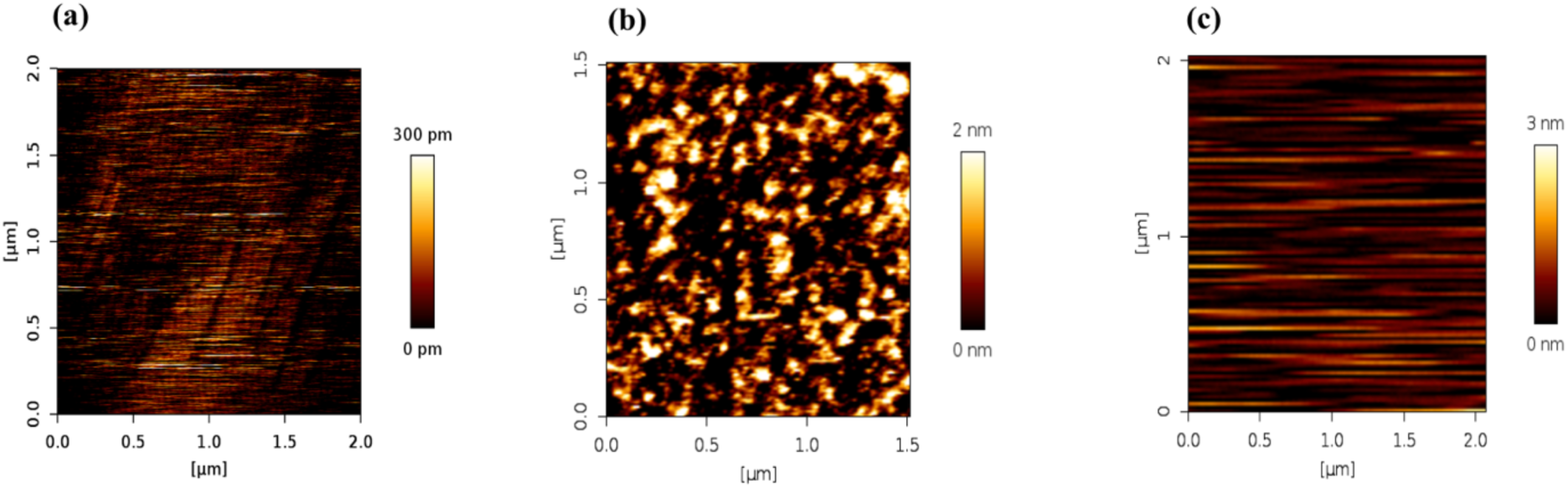
AFM images of (a) Supported Lipid Bilayer (SLB) before any protein treatment; (b) POPC:Chol 1:1 SLB after three rounds of buffer washes following pre-incubation with 100 nM MbxA; (c) POPC:Chol 1:1 SLB after three rounds of buffer washes following pre-incubation with 100 nM proMbxA.

### Sterol-induced membrane packing facilitates the insertion and oligomerization of acylated MbxA

To further elucidate the differences in lytic activity observed between MbxA and proMbxA, we examined the physical properties of membranes containing a variety of lipid compositions. We specifically focused on the fluidity and packing of the membrane, which can be measured by integrating the fluorescent dye Laurdan [42, 43]. Membrane fluidity was assessed by measuring the anisotropy, while membrane packing can be evaluated through general polarization (GP) measurements derived from the Laurdan emission spectrum across different lipid compositions. When a sterol such as cholesterol or ergosterol is incorporated into a membrane in a liquid-disordered state (T > Tm), it leads to closer packing of the membrane, resulting in a liquid-ordered state (Fig. 6a). Conversely, when a sterol is integrated into a membrane in a gel phase (T < Tm), it results in looser packing of the membrane, also yielding a liquid-ordered state (Fig. 6a).

**Figure 6:**
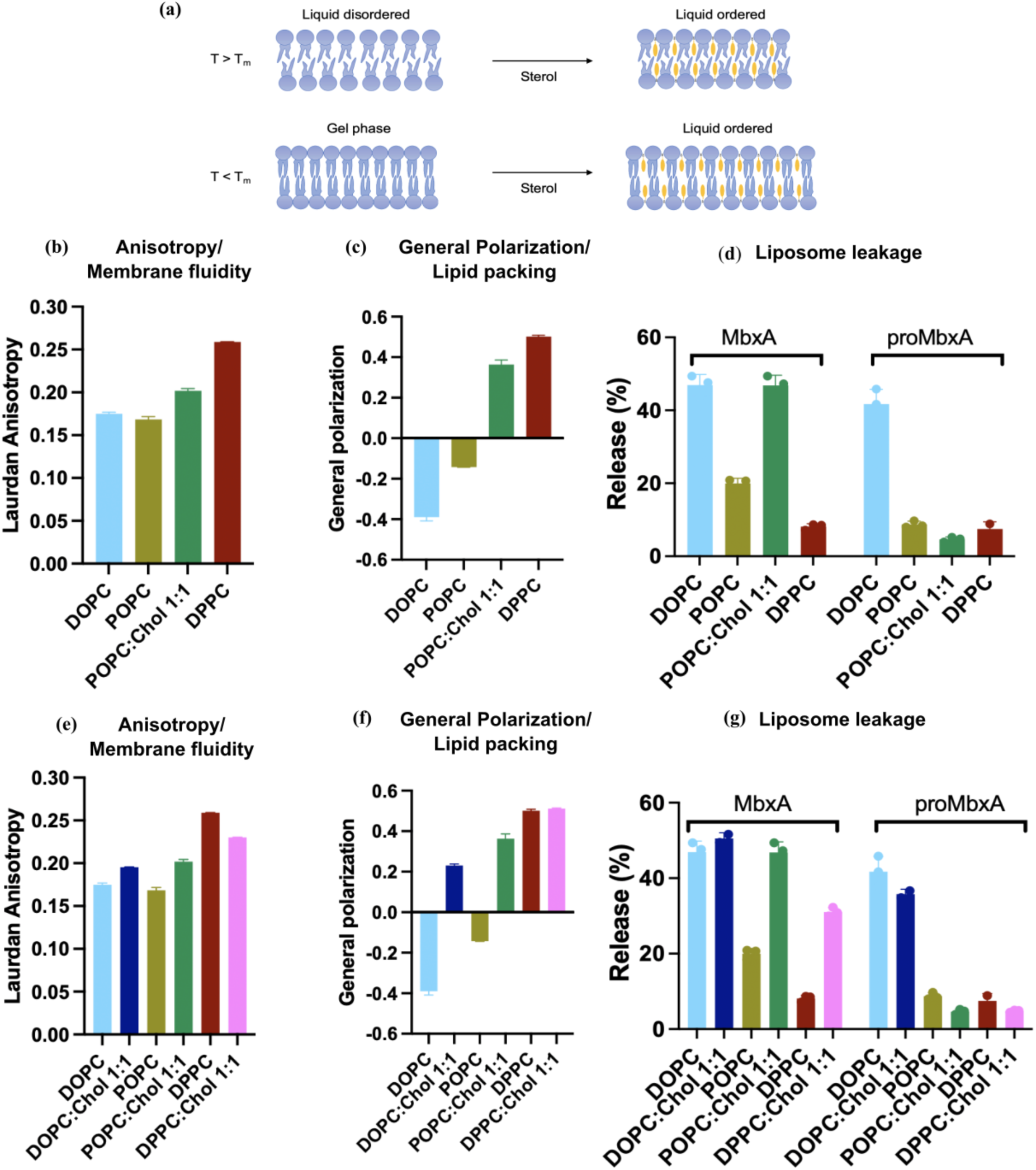
(a) Theory of lipid packing when sterol is introduced into a membrane in the liquid-disordered state and in the gel phase. (b, e) Anisotropy values of liposomes of different lipid compositions are calculated using the equation r = (Ivv - GIvh)/(Ivv + 2GIvh) where I is the fluorescence intensity; v and h are vertical and horizontal settings for the excitation and emission polarizers, respectively; G is the instrumental correction factor provided by the instrument for each instrument (further details are mentioned in the materials and methods section). A higher anisotropy measurement indicates less fluidic membranes and a lower anisotropy measurement indicates highly fluidic membranes. (c, f) General Polarization (GP) values calculated from the Laurdan emission spectrum using the equation (I435 - I500)/(I435 + I500), where I is the fluorescence intensity at the respective wavelengths mentioned. A higher GP value (more positive) indicates more closely packed membranes and a lower GP value (more negative) indicates loosely packed membranes. (d, g) Liposome leakage assay of MbxA and proMbxA on liposomes of different lipid compositions. (d) is a combined data of Fig. 2a and 2c. All the measurements are performed at 20°C. Lipid compositions with colour codes: **DOPC**, **DOPC:Chol 1:1**, **POPC**, **POPC:Chol 1:1**, **DPPC**, **DPPC:Chol 1:1**.

Anisotropy measurements indicate a similar degree of membrane fluidity for DOPC and POPC liposomes, while DPPC exhibits reduced fluidity (Fig. 6b). This is in accordance with previous observations [44], and indicates that the earlier observed difference in lytic activity between MbxA and proMbxA on POPC membranes (fig. 2a) is not a consequence of altered membrane fluidity. Concurrently, GP measurements reveal membrane packing in the order of DOPC < POPC < DPPC (Fig. 6c), which does align with the lytic activity of MbxA, observed in the order of DOPC > POPC > DPPC (Fig. 2a). In other words, MbxA demonstrates greater lytic activity when the packing of the membrane decreases.

Next, the effect of cholesterol on membrane packing and fluidity was examined. Since a POPC lipid membrane is in a liquid disordered state at 20°C (Tm; POPC=-2°C [45]), cholesterol incorporation strongly increased the GP of POPC membranes at 20°C resulting in membrane packing more similar to DPPC membranes (Fig. 6c). Likewise, the membrane fluidity of POPC liposomes decreased in the presence of cholesterol, indicated by an increased anisotropy (Fig. 6b). Based on the packing/fluidity results observed for membranes containing only PC lipids, a decrease in lytic activity of MbxA would be in accordance. However, the contrary is observed, as instead, cholesterol increases the lytic activity to levels comparable with DOPC membranes (Fig. 6d). As no such effect was observed for proMbxA, the specific role of the acyl-moieties in the lytic activity of this protein with cholesterol is highlighted once again.

To further specify the impact of cholesterol on membrane fluidity and packing, liposomes containing DOPC:Chol 1:1, and DPPC:Chol 1:1, were tested for anisotropy and GP (Fig. 6e,f) while assessing the lytic activity of MbxA and proMbxA on these formulations (Fig. 6g). In all cases, cholesterol incorporation stimulated the lytic activity of MbxA, although no such increase was observed for proMbxA (Fig. 6g). DOPC lipid membrane is in a liquid disordered state while DPPC membrane is in a gel phase at 20°C (Tm; DOPC=-17°C [46], DPPC=41°C [47]). Anisotropy measurements showed that cholesterol reduces the fluidity of DOPC liposomes, similar to POPC liposomes, while increasing the fluidity of DPPC liposomes (Fig. 6e). GP values for DOPC:Chol liposomes strongly increased, as observed for POPC:Chol liposomes, while the membrane fluidity of DPPC and DPPC:Chol 1:1 liposomes were similar, possibly caused by saturation of the signal (Fig. 6f). Altogether, these results clearly show the specific impact of cholesterol on the lytic activity of MbxA. Nevertheless, a secondary effect can be attributed to the membrane packing as an increase in GP coincides with reduced lytic activity both in the presence and absence of cholesterol.

Interestingly, while cholesterol incorporation into DOPC liposomes increased membrane packing (as indicated by GP measurements; Fig. 6d) and reduced membrane fluidity (as shown by anisotropy measurements; Fig. 6e) compared to DOPC liposomes, proMbxA was capable of inducing lytic activity toward DOPC:Chol 1:1 liposomes. This finding is in contrast with the results from POPC:Chol 1:1 and DPPC:chol 1:1 liposomes, where MbxA exhibited lytic activity, but proMbxA did not. This result indicates that the positive effect of cholesterol on the lytic activity of MbxA is not entirely dependent on the acyl-moieties of the protein. Previous studies have reported that incorporating cholesterol into DOPC membranes (but not for POPC or DPPC membranes) can lead to phase separation within the membrane [48–50]. If such phase separation occurs in DOPC:Chol 1:1 liposomes, it may facilitate the insertion of proMbxA into these phase-separated DOPC domains of the mixture, resulting in the release of liposomal contents, similar to that observed with DOPC liposomes.

### MbxA is unable to induce cytotoxicity when cholesterol is removed from human epithelial (HEp-2) cells

Although the above *in vitro* results clearly demonstrate the effect of cholesterol on the activity of MbxA, they hardly represent an *in vivo* membrane. Cellular membranes contain a wide variety of different lipids that vary in headgroup and lipid tail configuration. To investigate the *in vivo* effects of cholesterol on the cytotoxicity of MbxA and proMbxA, their lytic activity was tested on HEp-2 cells with and without cholesterol. HEp-2 cells contain 24.5% cholesterol [51]. To deplete cholesterol, varying concentrations of methyl beta-cyclodextrin (m-βCD) were used, after which the cholesterol content was assessed by GC-MS (Supplementary fig. 3a). As the viability of the cells decreased above 10mM m-βCD, a concentration of 7mM was chosen for the experiment. This reduces the total cholesterol content 12-fold, leaving approximately 2% of the total content of 24.5%. Cytotoxicity was assessed by observing plasma membrane permeabilization and membrane blebbing, which serve as indicators of MbxA-induced cytotoxicity, as described previously [34]. HEp-2 cells were stained with CellMask™ Deep Red to visualize the plasma membrane and suspended in a medium containing Sytox Green, a nuclear staining dye. Following incubation with MbxA, the plasma membrane of the HEp-2 cells became permeabilized, allowing Sytox Green to stain the nuclei (Fig. 7; first row). Additionally, MbxA cytotoxicity led to a membrane blebbing phenotype, with noticeable blebbing observed just 5 minutes after MbxA treatment (Fig. 7b, arrowheads). The size of the bleb and intensity of the nuclear staining slowly increases in the subsequent time points, which can be visualized in the 10-minute, 15-minute, and 20-minute images of MbxA treatment on cholesterol-intact HEp-2 cells (Fig. 7; first row). After 20 minutes of incubation, more pronounced nuclear staining by Sytox Green and an increase in the number of membrane blebs were evident. Supplementary Fig. 3b illustrates a broader area of HEp-2 cells subjected to MbxA, highlighting the Sytox Green permeabilization and membrane blebbing, while supplementary Fig. 3c shows the top layer of HEp-2 cells where membrane blebs are prominently visible. However, when cholesterol was removed from HEp-2 cells using 7 mM m-βCD, MbxA failed to induce cytotoxicity. No nuclear staining by Sytox Green or membrane blebbing was observed, even after 20 minutes of incubation with 100 nM MbxA (Fig. 7; second row). Notably, whether cholesterol was present or absent, proMbxA did not induce cytotoxicity in HEp-2 cells, consistent with previous observations [34] (Supplementary Fig. 3d).

**Figure 7:**
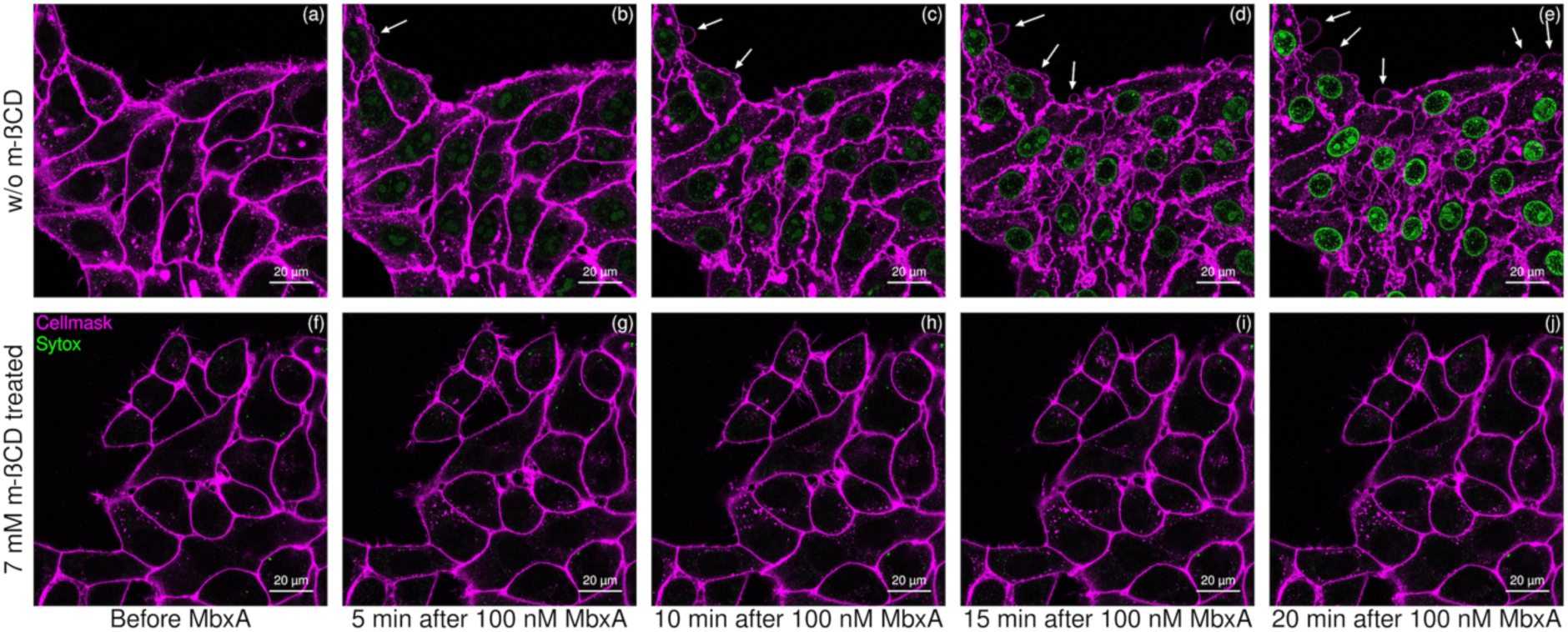
Confocal microscopy images of CellMask^TM^ Deep Red stained HEp-2 cells. First row (a,b,c,d,e): Nuclear staining of HEp-2 cells by Sytox green from the surrounding media due to the plasma membrane permeabilization and membrane blebbing (white arrowheads) resulting from the MbxA cytotoxicity in cholesterol- intact membranes of the HEp-2 cells. Second row (f,g,h,i,j): No nuclear staining nor membrane blebbing phenotype has been observed in HEp-2 cells when cholesterol-removed HEp-2 cells are incubated with 100 nM MbxA.

### MbxA & proMbxA bind to HEp-2 cell membranes with and without cholesterol

The flotation assay indicated that both MbxA and proMbxA can associate with POPC:Chol membranes, suggesting that the acyl chains are not required for membrane binding. To validate this finding *in vivo*, both MbxA and proMbxA were labeled with Atto-488, allowing for visualization of their binding to the HEp-2 cell membrane. HEp-2 cells were treated with 100 nM Atto488-MbxA^S9C^, and any unbound protein was subsequently washed away (Fig. 8b) to highlight the membrane-bound protein. As previously reported [34], with a lytic activity of 97% of the wild-type protein, Atto488-MbxA^S9C^ effectively bound to the plasma membrane of HEp-2 cells (Fig. 8c). Similar to MbxA, proMbxA was also observed to bind clearly to the plasma membrane of HEp-2 cells (Fig. 8i). A control experiment with 100 nM Atto-488 dye showed no binding to the HEp-2 cell membrane, further confirming that the binding of Atto488-MbxA^S9C^ and Atto488-proMbxA^S9C^ is due to the proteins themselves rather than free dye (Sup. Fig. 3). Additionally, a control experiment involving wheat germ agglutinin (WGA) binding to the HEp-2 cell membrane exhibited a binding pattern similar to that of Atto488- MbxA^S9C^ and Atto488-proMbxA^S9C^, reinforcing the conclusion that these proteins are indeed binding to the plasma membrane of the HEp-2 cells (Sup. Fig. 3).

**Figure 8:**
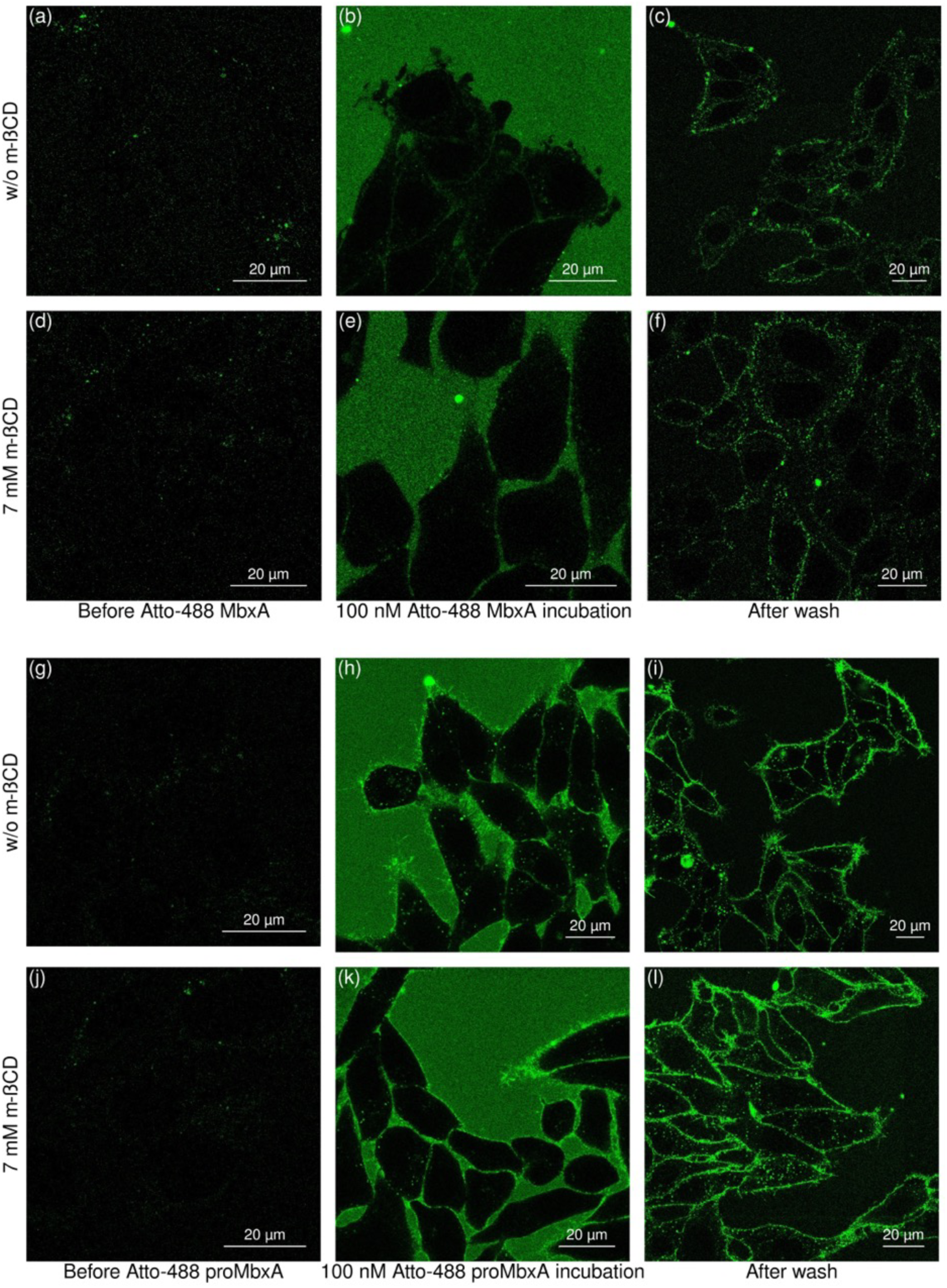
Confocal microscopy images of (a,b,c) cholesterol intact HEp-2 cells; (a) before incubating with Atto488-MbxA^S9C^, (b) incubation of HEp-2 cells with Atto488-MbxA^S9C^ which results in membrane blebbing, (c) HEp-2 cells after the excessive unbound protein being washed away. (d,e,f) Cholesterol removed HEp-2 cells using 7 mM m- βCD; (d) before incubating with Atto488-MbxA^S9C^, (e) incubation of HEp-2 cells with Atto488- MbxA^S9C^ which doesn’t show any membrane blebbing, (f) HEp-2 cells after the excessive unbound protein being washed away. (g,h,i) cholesterol intact HEp-2 cells; (g) before incubating with Atto488-proMbxA^S9C^, (h) incubation of HEp-2 cells with Atto488-proMbxA^S9C^ which doesn’t show any membrane blebbing, (i) HEp-2 cells after the excessive unbound protein being washed away. (j,k,l) Cholesterol removed HEp-2 cells using 7 mM m- βCD; (j) before incubating with Atto488-proMbxA^S9C^, (k) incubation of HEp-2 cells with Atto488-proMbxA^S9C^ which doesn’t show any membrane blebbing, (l) HEp-2 cells after the excessive unbound protein being washed away.

To investigate whether cholesterol in the host cell membrane is essential for the binding process of the toxin, Atto 488-labeled MbxA and proMbxA were added to both cholesterol-intact and cholesterol-removed HEp-2 cells (Fig. 8). Cholesterol-removed HEp-2 cells were incubated with both Atto488-MbxA^S9C^ (Fig. 8d, e, f) and Atto488-proMbxA^S9C^ (Fig. 8j, k, l), followed by washing with buffer to remove unbound proteins. Both MbxA^S9C^ (Fig. 8f) and proMbxA^S9C^ (Fig. 8l) were found to bind to the cholesterol-removed plasma membrane of HEp-2 cells, similarly to the binding observed in cholesterol-intact HEp-2 cells (Fig. 8a, b, c & Fig. 8g, h, i), indicating that cholesterol is not involved in the initial binding process of the protein to the membrane.

## Discussion

RTX toxins, such as MbxA from *M. bovis,* display virulence by disrupting the host cell membranes. Besides the β2 integrin receptor-dependent activity, RTX toxins can exhibit lytic activity via a receptor-independent interaction pathway, which is suggested to rely on specific interactions between the two acyl-moieties of the RTX toxin with the membrane [3, 23–33]. Here, we systematically characterized the role of the lipid membrane in the binding and lytic activity of MbxA (acylated at lysine residues K536 and K660 [34]), and its non-acylated counterpart proMbxA.

Introducing negatively charged lipids, exemplified by POPS (Fig. 2b), into neutral liposomes enhances the lytic activity of both MbxA and ProMbxA, illustrating this effect is independent on the acyl-moieties of the protein. This stimulating effect may be explained by a net positive surface charge density of the pore-forming hydrophobic domain at the N-terminus, as indicated by an model of proMbxA predicted by AlphaFold, which was modified according to SAXS data (Sup. Fig. 1). Putting this into context, negatively charged lipids are predominantly located in the inner leaflet of host cell membranes and therefore not accessible for RTX toxins [52]. However, during cell signaling or apoptosis, these lipids could be transiently exposed, potentially allowing the toxins to exploit these conditions for binding.

Similarly, introduction of unsaturated acyl chains in the liposomal lipids also increases the lytic activity of MbxA and Pro-MbxA (fig. 2a). This can be directly linked to changes in the biophysical properties of the membrane *i.e.*, decreased membrane packing and to a lesser extent increased fluidity (Fig. 6b,c), thereby facilitating the membrane its accessibility. Noteworthy, the lytic activity of proMbxA compared to MbxA is severely lower in POPC membranes, possibly highlighting a specific role for the acyl-chain moieties of the protein while interacting with the membrane. Although POPC membranes most closely resemble the acyl-chain configuration of native membranes [52], membrane packing and fluidity are also greatly influenced by sterols. Indeed, incorporating cholesterol (or ergosterol) into POPC liposomes significantly increased liposome leakage induced by MbxA, with leakage rising from 20% in POPC liposomes to 45% in POPC:Chol (1:1) liposomes (fig. 2c). On the contrary, a (small) decrease in lytic activity has been observed for proMbxA, clearly showing the importance of the protein its acyl-chain moieties. Remarkably, the presence of cholesterol deteriorated the earlier described favorable biophysical properties of the membrane. The membrane fluidity decreased, whereas membrane packing drastically increased to levels observed with DPPC liposomes. In other words, the enhanced lytic activity is caused by a specific interaction of acylated MbxA with cholesterol and not a result of an overall more accessible membrane.

Overall, the *in vitro* setup provides a systematic overview, displaying several factors (*e.g.,* lipid charge, membrane packing, sterols, etc.) that play a role in (pro)MbxA activity. Nevertheless, liposomes are simplified model membranes and do not necessarily represent the *in vivo* situation. Using HEp-2 cells as a model system, we tested (pro)MbxA activity in presence and absence of cholesterol. In short, MbxA only showed pore forming activity in normal cholesterol containing membranes, allowing the membrane impermeable Sytox Green to stain the nucleus. On the contrary, depleting the cholesterol content to 2% resulted in inactive MbxA. In the case of proMbxA, no lytic activity was observed, regardless of the cholesterol content, thereby confirming the earlier observed difference in the *in vitro* setup. Altogether, the *in vivo* experiments provide critical insights into the combined role of cholesterol and acylation in MbxA’s cytotoxic activity within a physiological context. The complete abrogation of lysis in cholesterol-depleted HEp-2 cells underscores its necessity for MbxA’s functional conformation and pore formation. This finding aligns with previous studies implicating the presence of cholesterol-rich regions in the activity of pore-forming toxins, further cementing cholesterol as a key lipid component for RTX toxin-mediated cytotoxicity [22, 29, 33, 53, 54].

A possible explanation for the cholesterol-dependent lytic activity of MbxA is that the presence of cholesterol results in the formation of more (stable) pores. To determine whether the acyl chains are essential for the binding of MbxA to cholesterol membranes, *in vivo* experiments using confocal microscopy were performed, showing both Atto-488-labeled MbxA^S9C^ and proMbxA^S9C^ were bound to the plasma membrane of HEp-2 cells, regardless of cholesterol presence (Fig. 8). Moreover, an *in vitro* flotation assay with liposomes revealed that after centrifugation both acylated MbxA and non-acylated proMbxA were present in the liposome fraction, although with slightly lower levels of proMbxA. These observations highlight that the initial membrane binding of MbxA is not dependent on its acyl-chains or on cholesterol. On the other hand, by AFM, it was observed that MbxA stayed tightly bound to the membrane, whereas proMbxA was removed upon washing, suggesting cholesterol may act as a stabilizer, enhancing pore stability and persistence in the lipid bilayer, likely through the protein’s acylation. The stabilizing role of cholesterol is consistent with findings from for example Herlax et al. (2009), who showed through FRET experiments that cholesterol enhances the oligomerization of HlyA in the host membrane [29].

Overall, these results highlight the intricate interplay between lipid composition and protein acylation on the pore-forming behavior of MbxA. However, the specific MbxA- membrane/cholesterol interactions at the molecular level remain unknown. There are several potential interpretations:

1. **Cholesterol-Mediated Membrane Organization**: Cholesterol may (re-)organize the local lipid environment, such that the insertion and oligomerization of acylated MbxA is favored. Such membrane restructuring could further potentiate the toxin’s lytic activity.
2. **Direct Acyl Chain-Cholesterol Interactions**: Alternatively, the acyl chains of MbxA might interact directly with cholesterol, aiding in the stabilization and clustering of the protein within cholesterol-enriched microdomains [25, 55, 56].
3. **Acylation-Induced Conformational Changes**: Finally, acylation may trigger conformational changes that expose specific cholesterol-binding motifs. In theory, exposure of these domains could stabilize membrane interactions and/or promote pore formation within the membrane, possibly by stimulating MbxA oligomerization. Sequence analysis of MbxA reveals the presence of 25 so-called CRAC/CARC motifs [57]. In fact, various other RTX proteins are known to possess these specific cholesterol recognition motifs, of which some are located within, or near, the pore-forming domain and found to directly interact with membrane cholesterol [22, 33, 53, 58].

In summary, our *in vitro* and *in vivo* studies elucidate the complex relationships between lipid composition, acylation, and the pore-forming mechanisms of MbxA. The findings highlight cholesterol’s essential role in enhancing MbxA’s lytic activity facilitated through protein’s acylation, even though the acyl chains and cholesterol are not needed for initial membrane binding. The varying effects of different lipid compositions show that secondary factors like membrane fluidity, packing, and surface charge also impact MbxA’s function, further emphasizing the complex interplay between protein and membrane. The specific interactions at the molecular level between acylated MbxA and cholesterol remain unknown, but could involve conformational changes that promote pore formation, possibly by activating cholesterol recognition motifs such as CRAC/CARC. The recognition of alternative sterols, like ergosterol, suggests some flexibility of MbxA in adapting to different lipid environments, thereby expanding its range of targets. Overall, these findings deepen our understanding of the molecular factors underlying RTX toxin activity and their role in membrane disruption and cytotoxicity.

## Materials and Methods

### Protein expression & purification

Expression, secretion, and purification of (pro)MbxA was performed as described before [34]. (pro)MbxA was expressed in BL21 (DE3) *E. coli* strains. The proteins were expressed in a two-plasmid system in which one plasmid carries the ABC transporter *hlyBD* gene, and the second carries either the *hlyC-mbxA gene* (for acylated MbxA) or *mbxA* gene (for proMbxA). The expressed proteins were secreted via HlyBD, culture supernatants were pooled, and purified by Immobilised Metal Affinity Chromatography (IMAC).

### Hemoglobin release Assay

Assays were performed as previously reported [34]. Defibrinated sheep blood cells were washed and centrifuged several times until supernatant became clear. Homogenized blood cells were incubated with 100nM of protein (MbxA, Atto488-MbxA^S9C^, proMbxA, Atto488- proMbxA^S9C^), for 30 minutes at 37°C. 16% SDS and IMAC buffer were used as positive (100%) and negative (0%) control. Cells were then centrifuged, supernatants analyzed for hemoglobin release, and quantified via OD on a FLUOstar OPTIMA microplate reader (BMG Labtech) at 544nm.

### Liposome preparation and leakage assay

Chloroform dissolved lipids were dried in a rotary evaporator at 40°C for 30 minutes. Lipids were resuspended in 1 mL ANTS buffer (12.5mM ANTS, 45mM DPX, 150mM NaCl, 20mM Tri-HCl pH 7.0). Lipid suspension was sonicated in six cycles (15s on/45s off) 50% duty cycle, and 50% (power) at RT. The resulting small unilamellar vesicles (SUVs) were flash-freezed in liquid nitrogen and thawed at room temperature (∼20 min) two times. Liposomes were extruded using a 100nm membrane pore. Untrapped ANTS+DPX were chromatographically removed through a sephedex G-50 column using 150mM NaCl, and 20mM Tris-HCl pH 7.0 (buffer A).

### GUV preparation and assay

Two indium tin oxide (ITO)-coated glass slides were cleaned with 70% ethanol and chloroform. Then lipid solution (2*10 μl) was spread on the ITO-coated side, and the lipid area of one slide was fenced by a ring of sigillum wax (VitrexTM). A chamber was prepared by pressing the ITO/lipid-coated sides to each other. The chamber was filled with 300μl of 10% sucrose solution and sealed. Incubated the chamber for 3h at RT applying an alternating current with 11Hz and 2V. GUVs were harvested using a GELoader® tip (Eppendorf). The chamber was incubated with BSA (1min) before adding GUVs to the observation chamber. The chamber was washed 3 times with PBS. Next, 250μL PBS + 50μL GUVs were added to the chamber. 100nM of (pro)MbxA was added and time-lapse images were captured for 30min with 30sec time interval.

### Atomic Force Microscopy

AFM was performed in liquid and imaging in intermittent contact mode (AC mode) in a JPK NanoWizard 3 atomic force microscope with NanoWizard Control Software v.5 version 5.0.84 by JPK (JPK, Berlin, Germany), using a non-conductive silicon nitride cantilever (DNP-S10- A, Bruker, Billerica, USA) with a nominal tip radius of 10 nm, a spring constant of 0.35 N/m and a resonance frequency of 65 kHz. Supported lipid bilayers were prepared using POPC or POPC:Cholesterol (1:1) liposomes. For this, 5µl of a 125µM stock was put on muscovite mica surface positioned inside a reservoir for liquid on a glass slide, and incubated with 5µl buffer (20mM Tris-HCl pH 7.0, 650mM NaCl) for 10min. 1ml of 50mM Tris-HCl pH 7.8, 400nM NaCl, and 10mM CaCl2 was added to the reservoir. 100nM of (pro)MbxA was added and incubated for 10min. Next, the mica was washed 3x with buffer. Again, 1ml of buffer was added and imaging was performed. All steps were performed at RT.

### Membrane lipid packing and membrane fluidity analysis

Membrane lipid packing was analyzed by measuring General Polarisation (GP) and membrane fluidity was analyzed by Anisotropy, by using a Horiba Fluorolog-3 fluorometer. 100μM SUVs (see liposome preparation section) were mixed with 0.3μM laurdan to achieve a dye:lipid ratio of 1:333. Samples were incubated at 37°C for 1h in the dark, shaking at 300rpm.

#### GP measurement

Emission spectra (400-600nm) were measured while exciting at 350nm (Entrance slit: 3.00 nm Bandpass; Exit slit: 3.00 nm Bandpass). GP calculation:

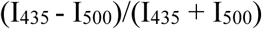

where I is fluorescence intensity.

#### Anisotropy measurement

Single point anisotropy measurement was performed by excitation at 350nm and an emission monitoring at 435nm. Anisotropy calculation:

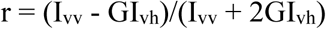

where I is fluorescence intensity; v (vertical), h (horizontal) settings for the excitation/emission polarizers respectively; G is the instrumental correction factor provided by the instrument.

### Flotation assay

SUVs (see liposome preparation section) were mixed with protein to a final volume of 100μL and incubated for 5min at RT. 100μL 60% sucrose solution was added, resulting in 30% sucrose solution with protein and liposomes. The mix was transferred into ultracentrifuge tubes of the AT3 rotor. 250μL of 20% sucrose solution was carefully added on top without disturbing the bottom layer. 50μL of liposome buffer was added on top and centrifuged at 80,000rpm for 1h at 4°C. After ultracentrifugation, samples were carefully taken from the three layers using a gel loading tip like follows: (1) 250μL - Bottom layer (2) 125μL - Middle layer (3) 125μL - Top layer. The same volume of 30% TCA (Trichloroacetic acid) was added to the three fractions. After incubation on ice (15min), samples were centrifuged (17,000xg; 15min; 4°C) and supernatant TCA removed. 500μL of ice-cold acetone was added to the pellet, resuspended, incubated on ice (15min) and centrifuged (17,000xg; 15min; 4°C). Supernatant acetone was removed, pellet dried using a speed vacuum centrifuge (10min). Pellet was resuspended in 30μL and analyzed by SDS PAGE.

### Cell culture and Live-cell imaging

HEp-2 cells were cultivated and passaged in DMEM (Pan Biotech) supplemented with 10% FCS, MEM vitamins, non-essential amino acids, amphotericin B (2.5ug/mL), and gentamicin (50ug/mL). Two days before, HEp-2 cells were seeded in 2mL DMEM in 35mm μ-Dish 1.5 H glass bottom dishes (Ibidi) and grown for 39-48 hours at 37°C under 5% CO2. Live-cell DMEM medium with 25mM HEPES in addition to the above-mentioned components is used for all the subsequent steps. Cells were washed with prewarmed DMEM live-cell medium and cell membranes stained using 5μg/mL CellMask Deep Red in DMEM-SG (containing 5μM Sytox Green), for 8min at 37°C and 5% CO2. Cells were washed with DMEM and replaced with 1 mL DMEM-SG. Images were acquired with Zeiss LSM880 Airyscan. 1mL DMEM-SG and 200nM protein was added into 1mL HEp-2 cells (100nM final). Images were captured every five minutes after incubation and activity was monitored as described before [34].

Atto488-labeled (pro)MbxA^S9C^ was used to visualize HEp-2 cell membrane binding. HEp-2 cells were washed with DMEM medium, replaced with 1mL of fresh DMEM medium, and imaged (Zeiss LSM880 Airyscan). Tile-scan images of 1mL HEp-2 cells were acquired prior to labeled (pro)MbxA and after 10 min incubation (100nM final) Unbound labeled proteins were washed off using DMEM and replaced with 1mL of fresh DMEM medium before image acquisition.

#### Microscopy settings

Confocal and Airyscan micrographs were recorded using a Zeiss LSM880 Airyscan microscope system (Carl Zeiss Microscopy GmbH) equipped with a Plan-Apochromat 63x/1.4 oil immersion objective lens. For excitation, a 405 nm Laser was used for Hoechst 33342, a 488 nm Argon laser for excitation of Sytox Green and Atto488-labeled MbxA^S9C,^ and a 633 nm laser for excitation of CellMask Deep Red.

### Mass spectrometry analysis of cholesterol removal

Cholesterol was removed from HEp-2 cells by incubation with m- βCD 5mM, 7mM, 10mM, 15mM, or 20mM (1h, 5% CO2). Prior to GC-MS analysis, 0.5 nanomol (nmol) of internal standard β-sitosterol was added. Lipids were extracted with 0.3mL n-butanol (2 times), solvent evaporated with nitrogen gas. Dried lipid films were resuspended in 100ul MSTFA and incubated (80°C; 30min).

1µl of derivatized compounds was injected and measured on a 5977B GC/MSD (Agilent Technologies) as described [59]. Oven temperature gradient: constant at 70°C for 1min, ramped at 42°C min^-1^ to 280°C, further with 4°C min^-1^ to 320°C, held constant for 3min (total 19min). Electron ionization source: 70eV; source temperature: 200°C, mass range: 60-600m/z, 5 scans per second. Metabolites were identified via MassHunter Qualitative (v b08.00, Agilent Technologies) by NIST14 Mass Spectral Library comparison (https://www.nist.gov/srd/nist-standard-reference-database-1a-v14). Authentic chemical standards for cholesterol were measured at a concentration of 5,10,50, 100µM and processed in parallel. Peaks were integrated using MassHunter Quantitative (v b08.00, Agilent Technologies). All metabolite peak areas were normalized to the sample amount (24K HEp-2 cells) and internal standard β-sitosterol (Sigma-Aldrich).

## Acknowledgments

We thank all members of the Institute of Biochemistry for fruitful and stimulating discussions. We are thankful for excellent technical support by Elisabeth Klemp and

## Data avaliability

All fluorescence micrographs will be uploaded tot he BioImage Database. All other data are avaliable from the corresponding author upon request.

## Supplementary figures

**Supplementary Figure 1:**
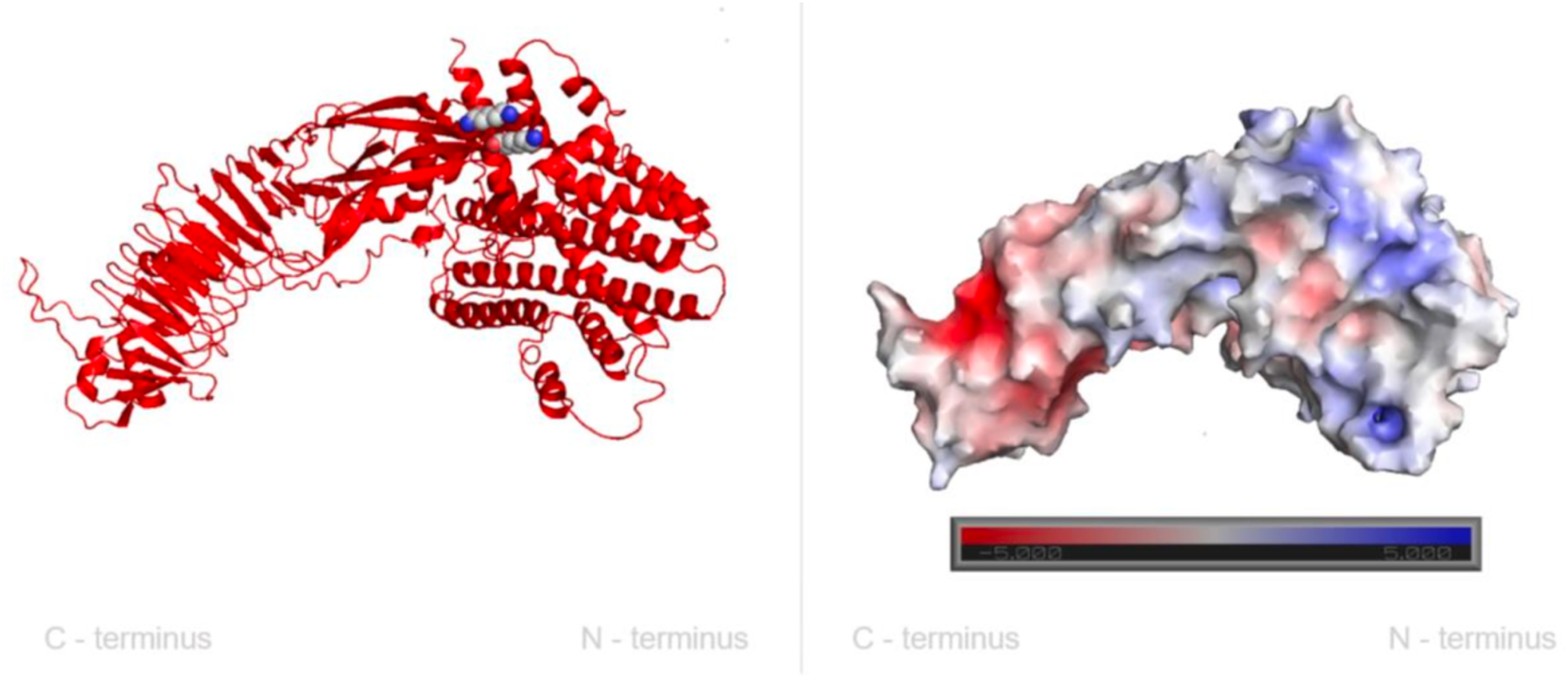
Left: AlphaFold predicted model of proMbxA, modified according to collected Small-Angle-X-ray-Scattering (SAXS) data. Right: Surface charge density representation of the model.

**Supplementary Figure 2:**
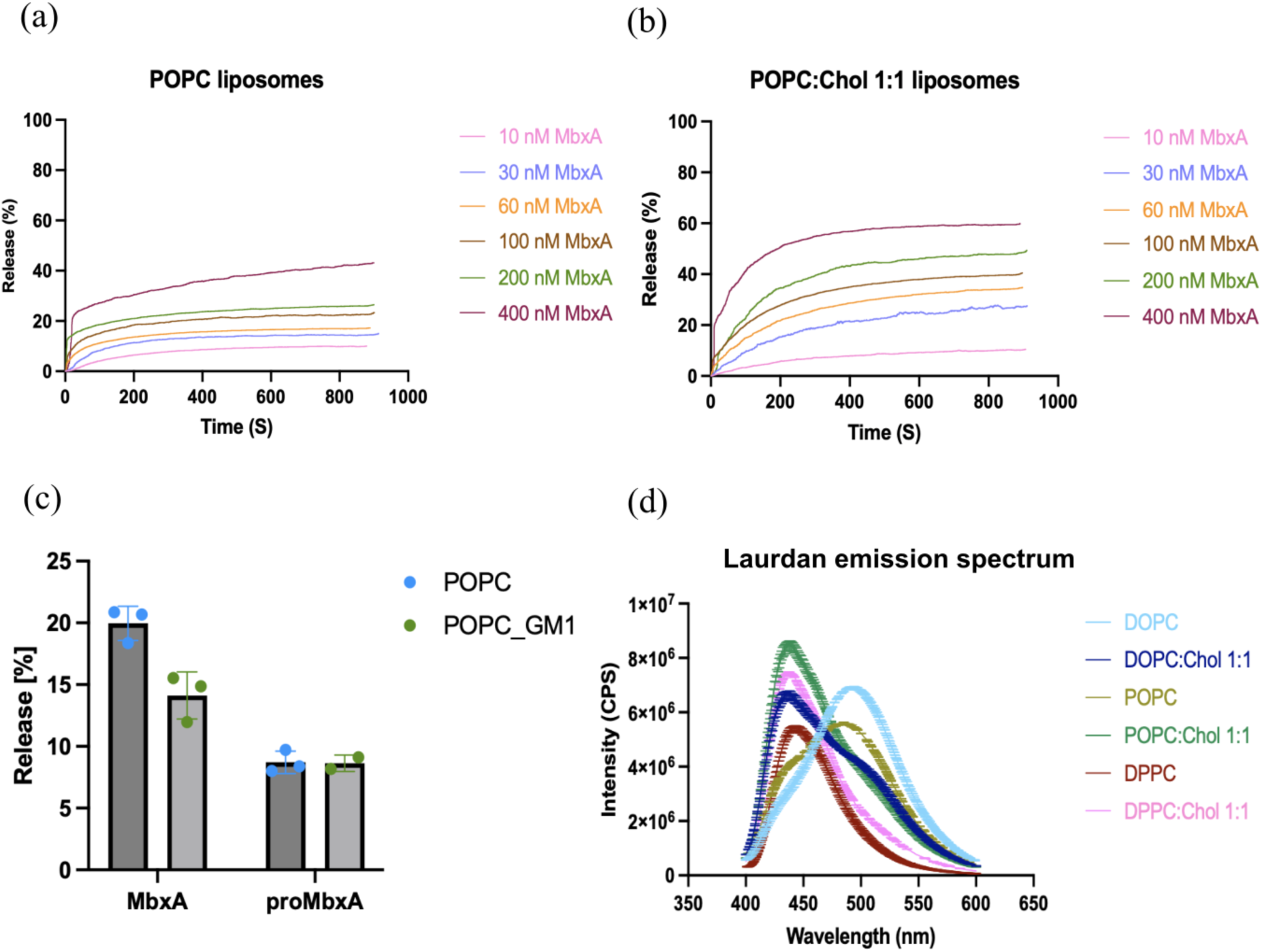
Panels (a&b) depict the time-based percentage release of liposomal content by the lytic activity of different MbxA concentrations of 10 nM, 30 nM, 60 nM, 100 nM, 200 nM & 400 nM on (a)12 μM POPC liposomes and (b) 12 μM POPC:Chol 1:1 liposomes. (c) The percentage of liposomal content released by the lytic activity of MbxA and proMbxA on 30% of GM1-incorporated liposomes. (d) Laurdan emission spectrum of liposomes of different lipid compositions from 400 nm to 600 nm at an excitation wavelength of 350 nm.

**Supplementary Figure 3:**
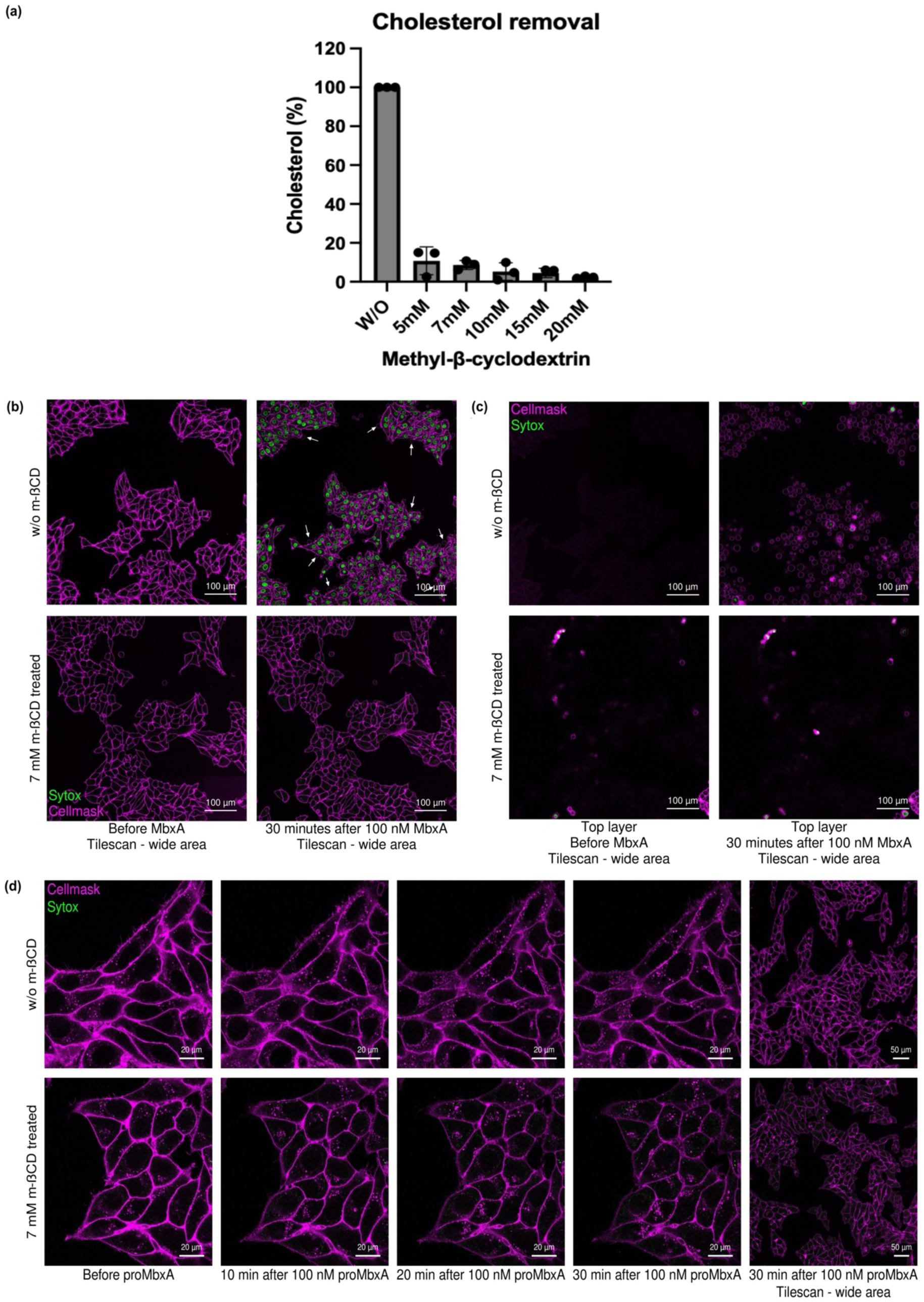
(a) Mass spectrometry data of cholesterol content in the HEp-2 cells when cholesterol was removed from HEp-2 cells using different concentrations of methyl beta-cyclodextrin. (b) A wider area of HEp-2 cells showing membrane permeabilization and membrane blebbing phenotypes when cholesterol is intact (First row) and the absence of MbxA cytotoxicity when cholesterol is removed from HEp-2 cells; therefore there is no membrane permeabilization nor membrane blebbing (Second row) when the cells are treated with 100 nM MbxA. (c) A layer above the HEp-2 cells provides a detailed view of the membrane blebs resulting from MbxA cytotoxicity when cholesterol is intact in HEp-2 cells (Upper row). At the same time, cholesterol-removed HEp-2 cells don’t show membrane blebs upon MbxA incubation when cholesterol is removed from the HEp-2 cells (Lower row). (d) When HEp-2 cells are incubated with 100 nM proMbxA, both cholesterol-intact (Upper row) and cholesterol-removed (Lower row) HEp-2 cells were unable to induce membrane permeabilization nor membrane blebbing phenotypes.

**Supplementary Figure 4:**
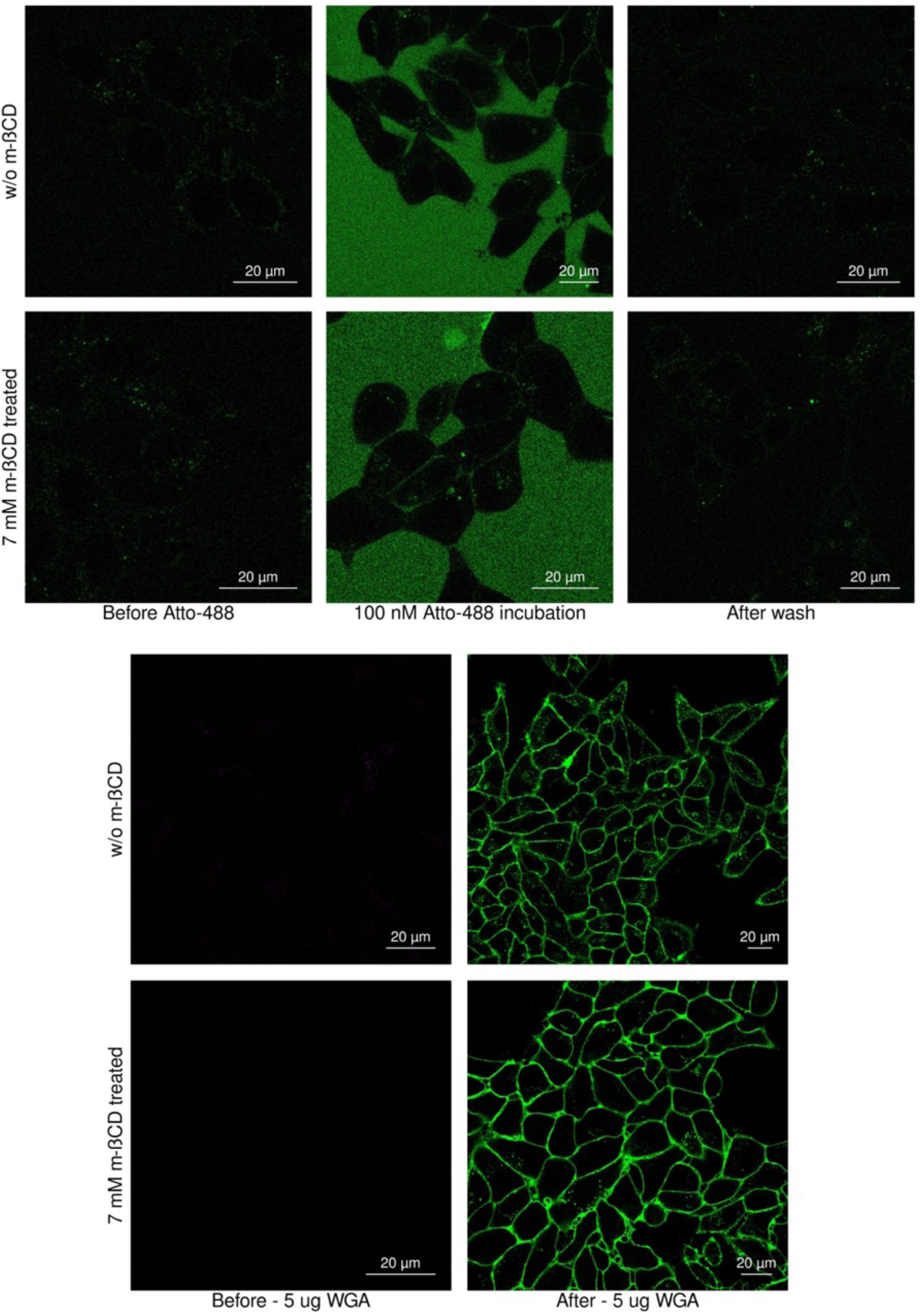
Control experiments by incubating 100 nM Atto-488 dye on (First row) cholesterol intact HEp-2 cells and (Second row) cholesterol removed HEp-2 cells; (left) before incubation, (middle) during incubation, (right) after washing excess dye using buffer. Control experiments by incubating 100 nM Wheat Germ Agglutinin (WGA) on (third row) cholesterol intact HEp-2 cells and (fourth row) cholesterol removed HEp-2 cells; (left) before incubation (right) 10 minutes after incubation.

**Supplementary Figure 5:**
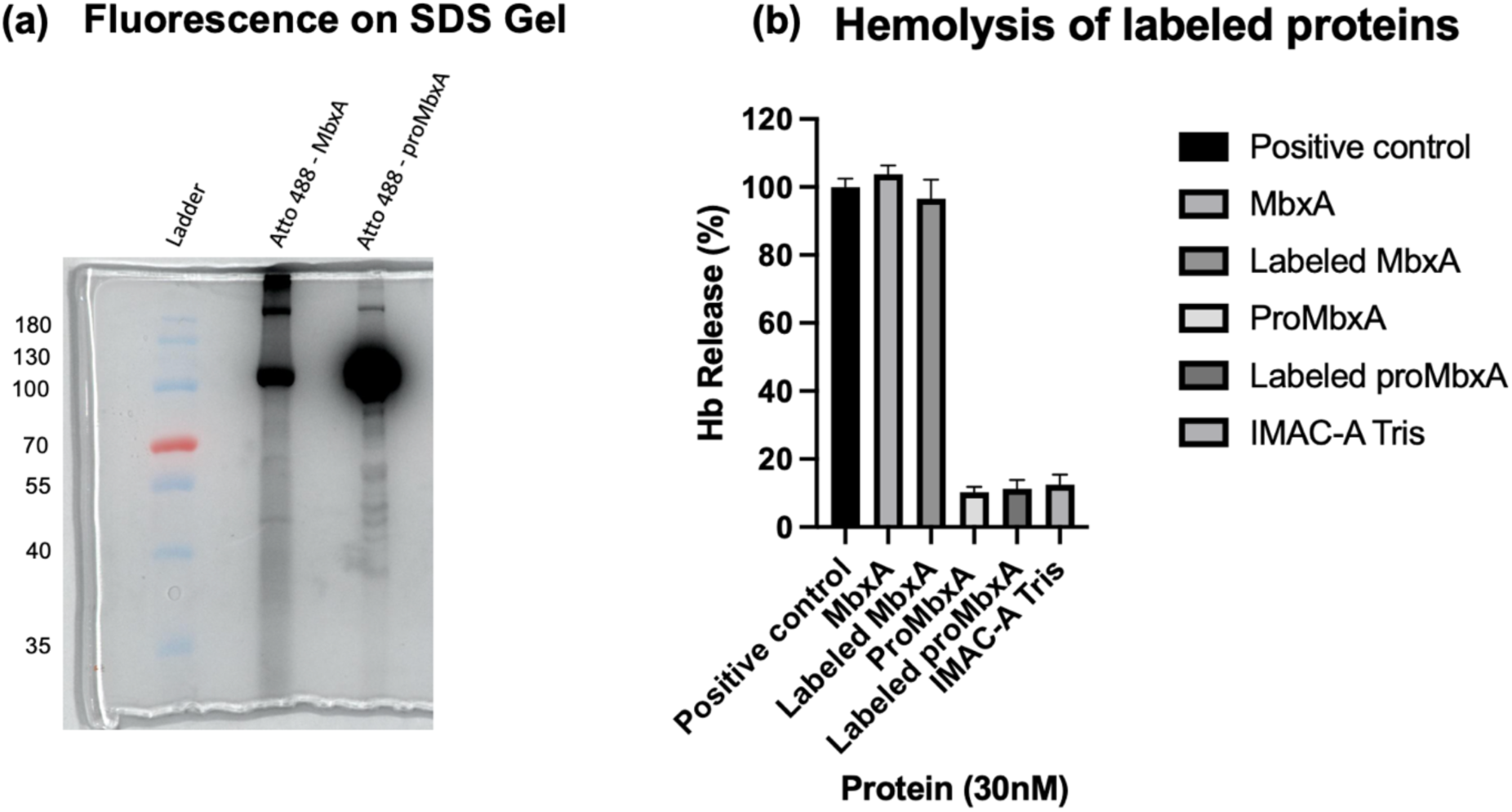
(a) 10% SDS gel showing the fluorescence of Atto-488-labeled MbxA^S9C^ and proMbxA^S9C^. (b) Hemoglobin release assay showing the activity of proteins.

